# A multi-tissue *de novo* transcriptome assembly and relative gene expression of the vulnerable freshwater salmonid *Thymallus ligericus*

**DOI:** 10.1101/2024.03.21.586089

**Authors:** Giulia Secci-Petretto, Steven Weiss, André Gomes-dos-Santos, Henri Persat, André M. Machado, Inês Vasconcelos, L. Filipe C. Castro, Elsa Froufe

**Affiliations:** CIIMAR/CIMAR, Interdisciplinary Centre of Marine and Environmental Research, University of Porto, Terminal de Cruzeiros do Porto de Leixões, Avenida General Norton de Matos, S/N, 4450-208 Matosinhos, Portugal; Department of Biology, Faculty of Sciences, U. Porto - University of Porto, Portugal; University of Graz, Institute of Biology, Universitätsplatz 2, A-8010 Graz, Austria; Société Française d’Ichthyologie, Muséum National d’Histoire Naturelle Paris, France, 57 rue Cuvier CP26, 75005 Paris, France

**Keywords:** *Thymallus*, transcriptome assembly, relative gene expression, Salmonidae

## Abstract

Freshwater ecosystems are among the most endangered worldwide. While numerous taxa are on the verge of extinction as a result of global changes and direct or indirect anthropogenic activity, genomic and transcriptomic resources are a key tool to comprehend species’ adaptability and serve as the foundation for conservation initiatives. The Loire grayling, *Thymallus ligericus*, is a freshwater European salmonid endemic to the upper Loire River basin. It comprises fragmented populations dispersed over a small area and has been identified as a vulnerable species. Here, we provide a multi-tissue *de novo* transcriptome assembly of *T. ligericus*. The final assembly, with a contig N50 of 1,221 and Benchmarking Universal Single-Copy Orthologs (BUSCO) scores above 94%, is made accessible, along with structural and functional annotations and relative gene expression of five tissues. This is the first transcriptomic resource for this species, and it will serve as the foundation for future research on a species that is increasingly exposed to environmental stressors.

## Introduction

Freshwater fishes are among the most threatened vertebrates. In Europe for example, they are the second most threatened group of organisms following molluscs (Costa et al. 2021). Of the 434 fish species reported in the IUCN red list, an estimated 41.2% are threatened, salmonids being the second most affected family (Costa et al. 2021). The most common threats affecting freshwater fauna in Europe include habitat fragmentation, pollution, the presence of invasive species, dams and hydropower plants and climate change (Mueller et al. 2020). Thymallinae (graylings), a monogeneric subfamily in the diverse Salmonidae family, is represented solely by the genus *Thymallus*, commonly named grayling. According to the latest revision, 15 taxa are currently listed in the genus (Weiss et al. 2021). The highest diversity occurs in the Asian continent, where 12 of the 15 known species are found (Weiss et al. 2021). One of these, *Thymallus arcticus* (Pallas, 1776), also inhabits some parts of North America, more specifically Alaska, Montana and Canada (Weiss et al. 2021). Three species are recognized in Europe: the widely distributed *Thymallus thymallus* (Linnaeus, 1758), the Adriatic grayling *Thymallus aeliani* Valenciennes, 1848 and the Loire grayling *Thymallus ligericus* Persat, Weiss, Froufe, Secci-Petretto & Denys, 2019. Until a few years ago, the latter two were considered different lineages of *T. thymallus. Thymallus ligericus* is the most recently described *Thymallus* species (Persat et al. 2019) and a revision of the genus categorized it as vulnerable, based on its distribution range and the documented habitat fragmentation (Persat et al. 2016; Weiss et al. 2021). This grayling inhabits the upper reaches of the Loire River (France) and some of its main tributaries, such as the Vienne, the Allier, the Sioule and the Lignon Rivers (Persat et al. 2016; Weiss et al. 2021). Populations of *T. ligericus* have maintained their reproductive isolation despite decades of failed restocking attempts as demonstrated by the *T. ligericus* population in the Vienne River, which has been confronted with hatchery-reared *T. thymallus* over the course of 50 years, yet no trace of these stocked individuals can be found (Persat et al. 2016).

The impact of environmental changes on species functions such as physiology, behaviour, and gene expression, is relatively unexplored in Thymallinae. Studies exist for the widely distributed *T. thymallus* and *T. arcticus*, as well as for other salmonid species and can offer insights into how environmental stressors may affect *T. ligericus* and other Thymallinae. Indeed it is demonstrated that events such as rising temperature, discharge fluctuations and human pollution affect physiological functions in *T. thymallus* and *T. arcticus*, as well as *Salmo trutta* Linnaeus, 1758 *and Salmo salar* Linnaeus, 1758 e.g. by inducing male bias, disrupting reproduction, growth, recruitment and feeding and swimming behaviours (Newcombe and Jensen 1996; Deegan et al. 1999; Luecke and MacKinnon 2008; Wedekind et al. 2013; Richard et al. 2015; Warren et al. 2015; Bašić et al. 2018; Opinion et al. 2020; Hayes et al. 2021; Auer et al. 2022). Thymallinae are strictly freshwater salmonids and their migratory behaviour within the same watercourse during the spawning season (West et al. 1992; Hughes and Reynolds 1994; Meyer 2001; Heim et al. 2015; Zuev et al. 2021) make them particularly sensitive to stream disturbances. The Loire is one of Europe’s longest rivers. Its first largest tributary, the Allier River, includes several biodiversity hotspots for plants and birds (Moatar et al. 2022) and hosts populations of *T. ligericus*. Some major urbanization of the Loire basin has arisen in the Allier Valley, impacting the environment with anthropogenic activities such as the construction of dams (Moatar et al. 2022). In 1994 the implementation of the “Loire Grandeur Nature Plan” gradually began to restore and improve the management of the Loire and Allier river environments, which had suffered from unstopped anthropogenic activities for decades (Moatar et al. 2022). Despite this, effects of climate change have been visible in this area and the Allier River, an important regional resource in terms of domestic and irrigation water supply, has recently experienced extreme dry summers with insufficient winter recharge (Labbe et al. 2023). These extreme conditions have inevitable consequences on the ecosystem’s biota and developing mechanisms to cope with such environmental fluctuations is critical for organisms’ survival. Understanding the limits of species resilience is vital to ensure the preservation and correct management of ecosystems. Genomic and transcriptomic resources, coupled with studies of ecology and physiology, are key tools that can help uncover compensatory mechanisms such as gene expression regulation in extreme environmental conditions. Increasingly lower sequencing costs coupled with advances in high-throughput sequencing techniques allow the generation of genomic and transcriptomic resources for non-model species (Connon et al. 2018; Gomes-dos-Santos et al. 2022). Among Thymallinae, transcriptomic resources have only been generated for *T. thymallus* (Pasquier et al. 2016; Varadharajan et al. 2018), which is also the only species of the genus with an available whole genome assembly (Varadharajan et al. 2018; Savilammi et al. 2019). Given the critical conservation status of *T. ligericus*, the lack of transcriptomic resources for this species may hinder future efforts to prevent its decline. In this work, we generate a multi-tissue transcriptome of the vulnerable species *Thymallus ligericus* from the Alagnon River (Allier basin). Additionally, we provide relative gene expression profiles for five tissues. As no transcriptomic resources are yet available for this taxa, this work provides a valuable framework for future studies that will assess the species’ ability to respond to stressors such as climate change and anthropogenic activity, to which *T. ligericus* is constantly exposed.

## Methods

### Animal Sampling

At the end of the summer season (27/08/2022), one adult male of *T. ligericus* was caught by angling from the Alagnon River (45° 14’ 34.4 N-3° 10’ 23” E; Fig. 1) in the Massiac district (Loire, France). Afterwards, 1cm^3^ of tissue was sampled from the brain, gills, gonad, liver, and muscle. Tissues were then stored in RNA-later at 4ºC for 24 hours and later at -80°C in preparation for RNA extraction. The samples were collected with the permission of the Direction Départementale des Territoires-Préfète de la Loire and genetic analyses were carried out according to the Nagoya Protocol (reference permit number: TREL2206915S/602).

**Fig. 1.**
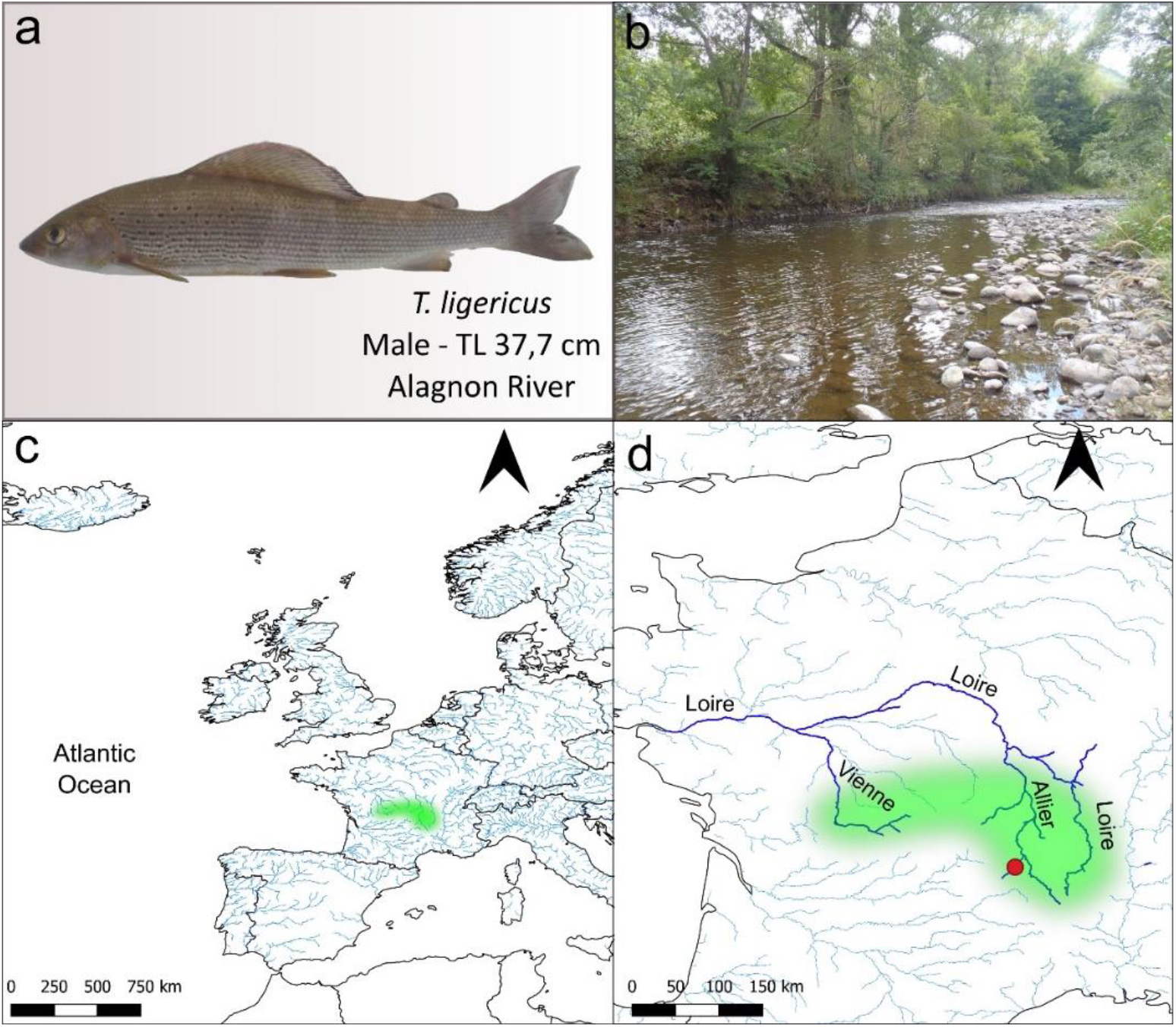
Specimen analyzed and detailed representation of its area of occurrence. a) Male *T. ligericus* analyzed in the present work. TL: total length. b) Photo of the Alagnon River section where the specimen was caught. c) Map with approximate distribution (marked in green) of *T. ligericus* based on available literature^3-5^. b) Zoom in on the species distribution in the Upper Loire River. The red dot identifies the sampling point (Alagnon River) where the specimen was caught.

### RNA extraction, sequencing and library construction

Total RNA was extracted from the five tissues with NZY Total RNA isolation kit (NZYtech) following the manufacturer’s instructions. RNA concentration was measured with DeNovix DS-11 FS and its integrity was assessed by running 3 μl in 1% agarose gel. The five samples were then sent in dry ice to Macrogen facilities (Seoul, South Korea) where strand-specific libraries with 250-300 bp insert sizes were built. Samples were sequenced using 150 bp paired-end reads on a Novaseq 6000 platform.

### Raw data quality control and assembly

The quality of the raw reads was assessed for each sample with FastQC v0.11.8 (Andrews, 2010). The software Trimmomatic v0.38 (Bolger et al. 2014) was then used to quality-filter the reads and trim the Illumina adapters with the following parameters: LEADING-5 TRAILING-5 SLIDINGWINDOW-5:20 MINLEN-36. Afterwards, random sequencing errors were fixed with the program Rcorrector v1.0.3 Song and Florea, 2015).

With no *T. ligericus* genome currently available, we opted to assemble the transcriptome with a *de novo* approach. Assembly was carried out with Trinity v2.13.2 (Grabherr et al. 2011; Haas et al. 2013) with default parameters. Afterwards, transcript redundancy was removed with Corset v1.0.9 (Davidson and Oshlack, 2014). To remove possible contamination, the assembled transcripts were then matched against the NCBI nucleotide database NCBI-nt; (Download; 24/08/2021) and Univec database (Download; 02/04/2019) in Blast-n v2.11.0 (Camacho et al. 2009). Transcripts with less than 100 bp, an e-value of 1e-5, an identity score <90% and with no match to the class Actinopterygii (Taxonomy ID:7898) were removed from the assembly, as well as everything that matched against the Univec database. The completeness of the transcriptome was assessed both before and after redundancy/contamination removal with BUSCO v3.0.2 (Simão et al. 2015) with lineage-specific libraries of Eukaryota, Metazoa, Vertebrata and Actinopterygii. Transrate v1.0.3 (Smith-Unna et al. 2016) was then used to assess transcriptome integrity before and after redundancy/contamination removal.

### ORF prediction and transcriptome annotation

Open Reading Frames (ORFs) prediction was carried out with Transdecoder v5.3.0 (https://transdecoder.github.io/). The software Blast-p v2.12.0 (Camacho et al. 2009) and hmmscan of hmmer2 package v2.4 (Finn et al. 2011) were used to search for homology and proteins by matching the transcripts against UniProtKB/Swiss-Prot48 (Bateman et al. 2017) and PFAM (Punta et al. 2012) databases. Both TransDecoder output in .gff format and the *de novo* assembly were processed with Gtf/Gff Analysis Toolkit (AGAT) v0.8.0 (Dainat et al. 2020) to prepare the file .gff3 for functional annotation. Afterwards, AGAT tool was used to standardize file formatting of protein and transcript fasta files. Functional annotation was carried out with InterProScan v5.44.80 (Quevillon et al. 2005) and Blast-n/p/x search. The proteins were mapped against NCBI-RefSeq – Reference Sequence Database (Download; 10/03/2022), NCBI-nr – non-redundant database (Download; 15/12/2021) and InterPro database (Download; 30/03/2019) using Blast-p/x tools implemented in DIAMOND v2.0.13 (Buchfink et al. 2014). The transcripts were searched in nt and nr databases of NCBI using Blast-n and Blast-x tools, respectively, from NCBI and DIAMOND. Finally, the blast outputs and InterProScan outputs were concatenated and integrated into the annotated gff3 file using AGAT. Gene names were assigned following the best blast hit and ranked as follows: 1 for Blast-p Hit from RefSeq database; 2 for Blast-p Hit from NCBI-nr database; 3 for Blast-x Hit from NCBI-nr database; 4 for Blast-n Hit from NCBI-nt database.

### Relative gene expression

Relative gene expression was assessed in each of the five tissues using two distinct methods: the first approach involved utilizing the *de novo* assembled transcriptome. This analysis was conducted with Trinity scripts, aligning the trimmed and corrected reads against the transcriptome using Bowtie2 v2.3.0 (Langmead and Salzberg, 2015) and quantifying gene expression with RSEM v1.2.31 (Li and Dewey, 2014). For the second approach, a reference genome-based pipeline was applied, using the available *T. thymallus* genome as a reference. The reads were mapped to the *T. thymallus* genome using HISAT2 v2.2.1 (Kim et al. 2015), and the resulting SAM files were converted to BAM format and sorted using SAMtools v1.9 (Li et al. 2009). StringTie v2.1.2 Pertea et al. 2015) was used to generate read counts for each mapped dataset.

## Results

### RNA sequencing

Raw reads obtained for each tissue were deposited at the NCBI Sequence Read Archive database under the accession numbers SRR26130661-SRR26130665 and the BioProject PRJNA1019285. The non-redundant transcriptome assembly is provided on NCBI databases under the accession number GKPV00000000. Both redundant and non-redundant transcriptome assemblies (^*^_trinity_filtered.fasta.gz and ^*^_transcriptome.fasta.gz) are also provided on Figshare platform (doi: 10.6084/m9.figshare.24174285), together with transcripts (^*^_genes_final.fasta.gz), mRNA (^*^_mrna_final.fasta.gz), ORF prediction (^*^_cds_final.fasta.gz), proteins ORF prediction (^*^_proteins_final.fasta.gz), annotation files (*_annotation.gff3.gz) and relative gene expression results.

### Raw datasets and pre-assembly processing quality control

The sequencing produced 18,708,599 paired-end raw reads for the brain tissue, 22,621,561 for the gill, 23,200,311 for the gonad, 23,250,910 for the liver and 23,118,988 for the muscle (Table 1).

**Table 1.** Number of raw reads obtained for each of the five tissues and number of reads retained for the assembly after Trimmomatic removal.

|  | Brain | Gill | Gonad | Liver | Muscle |
| --- | --- | --- | --- | --- | --- |
| Raw sequencing reads | 18,708,599 | 22,621,561 | 23,200,311 | 23,250,910 | 23,118,988 |
| Reads after Trimmomatic clean-up | 18,285,939 | 22,130,039 | 22,718,756 | 22,742,055 | 22,682,786 |

After read trimming and correction (carried out respectively with Trimmomatic and Rcorrector) the quality check performed in fastQC returned high Phred score values (>20) for both forward and reverse sequences (Fig. 2). The number of reads retained after trimming and used in the assembly are shown in Table 1.

**Fig. 2.**
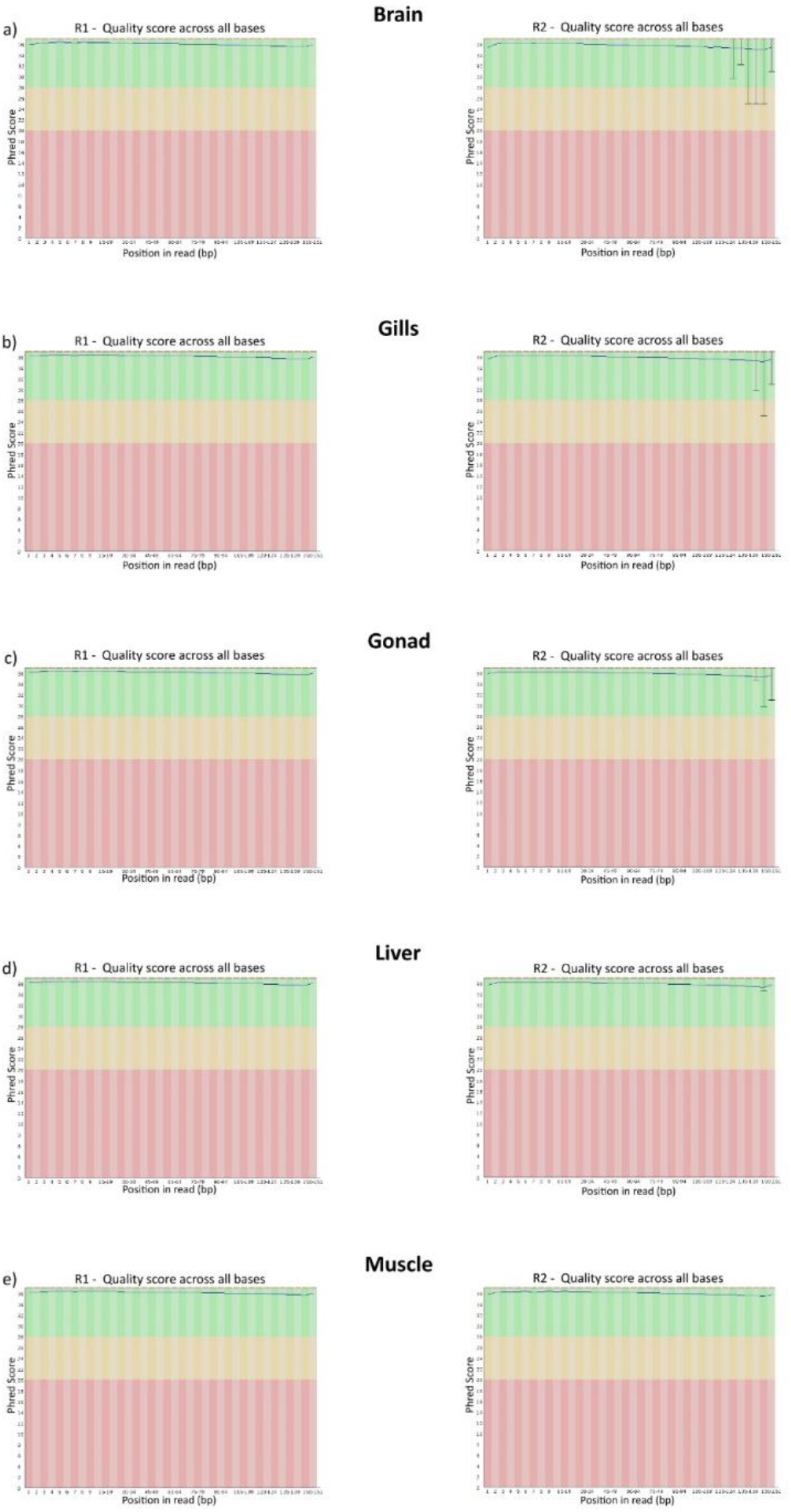
Quality report of the trimmed reads obtained in FastQC for each of the five tissues. a) brain; b) gills; c) gonad; d) liver; e) muscle.

### Transcriptome assembly, annotation and relative gene expression

The newly assembled redundant transcriptome included 1,147,286 sequences and an N50 length of 922 (Table 2). The Benchmarking Universal Single-Copy Orthologs (BUSCO) allowed evaluation of the gene completeness by matching the transcriptome with Eukaryota, Metazoa, Vertebrata and Actinopterygii orthologues database (respectively 303, 978, 2,586 and 4,584). BUSCO scores show a high level of gene completeness with more than 90% (complete + duplicated) genes for the four inferred groups (Euk, Met, Vet, Act) (Table 2).

**Table 2.** Transrate and BUSCO scores of the redundant and non-redundant transcriptome assembly. Euk: Eukaryota; Met: Metazoa; Vet: Vertebrata; Act: Actinopterygii. T: Total BUSCOs found complete + fragmented); C: complete BUSCOs; (single + duplicated); S: Complete and single-copy BUSCOs; D: Complete and duplicated BUSCOs; F: Fragmented BUSCOs; M: Missing USCOs.

| Transrate scores |  |  |  |  |  |  |  |  |  |  |  |
| --- | --- | --- | --- | --- | --- | --- | --- | --- | --- | --- | --- |
|  | Number of transcripts | N bases | Mean transcript length (bp) | Number of transcripts over 1 K nt | Number of transcripts over 10 K nt | N90 transcript length (bp) | N70 transcript length (bp) | N50 transcript length (bp) | N30 transcript length (bp) | N10 transcript length (bp) | Percentage of GC (%) |
| Redundant | 1,147,286 | 752,159,332 | 655.58151 | 177,319 | 428 | 282 | 492 | 922 | 1,755 | 3,620 | 0.44874 |
| Non-redundant | 279,365 | 280,214,152 | 1003.03958 | 90,928 | 113 | 509 | 823 | 1,221 | 1,909 | 3,529 | 0.44524 |
| Busco analysis (%) |  |  |  |  |  |  |  |  |  |  |  |
|  | Euk (n: 303) |  | Met (n: 978) |  | Vet (n: 2,586) |  | Act (n: 4,584) |  |  |  |  |
| Redundant | T:100;C:99.0%[S:20.8%,D:78.2%],<br>F:1.0%,M:0.0% |  | T:100;C:99.2%[S:22.1%,D:77.1%],<br>F:0.8%,M:0.0% |  | T:98.2;C:89.7%[S:24.0%,D:65.7%],<br>F:8.5%,M:1.8% |  | T:94;C:84.6%[S:23.5%,D:61.1%],<br>F:9.4%,M:6.0% |  |  |  |  |
| Non-redundant | T:99.4;C:99.1%[S:74.3%,D:24.8%],<br>F:0.3%,M:0.6% |  | T:98.2;C:97.4%[S:71.0%,D:26.4%],<br>F:2.0%,M:0.6% |  | T:97.6;C:88.7%[S:72.2%,D:16.5%],<br>F:8.9%,M:2.4% |  | T:92.6;C:83.3%[S:65.7%,D:17.6%],<br>F:9.3%,M:7.4% |  |  |  |  |

After the assembly, redundancy removal was carried out with Corset. The program removed about 75% from the initial amount of transcripts increasing the N50 from 922 to 1,221 and causing a general improvement in the quality of the assembly (Table 2). This is also reflected in the BUSCO scores. Although the percentages of matched genes are slightly lower, the percentages of single-copy genes increased significantly in the four classes (Euk, Met, Vet, Act) (Table 2), highlighting the importance and efficiency of redundancy removal. Afterwards, the non-redundant transcriptome was decontaminated with Blast-n searches against the NCBI-nt and Univec databases. Of the 68,242 predicted proteins 54,176 were functionally annotated (Table 3). All the annotation files can be consulted in FigShare (doi: 10.6084/m9.figshare.24174285).

**Table 3.** Statistics from the structural and functional annotation of the final transcriptome assembly.

| Structural annotation | N transcripts | N cdss | N exons | Tot gene length | Tot cds length | Tot exons length | Mean gene length | Mean cds length | Mean exons length |  |
| --- | --- | --- | --- | --- | --- | --- | --- | --- | --- | --- |
|  | 279,442 | 68,242 | 68,242 | 280,286,098 | 57,054,459 | 110,810,116 | 1,003 | 836 | 1,623 | - |
| Functional annotation<br>InterPro/Blast* | CDD | Coils | GO | Gene3D | Hamap | InterPro | KEGG | MetaCyc | MobiDBLite | PIRSF |
|  | 15,539 | 9,158 | 26,321 | 33,102 | 384 | 41,633 | 1,865 | 1,565 | 27,711 | 1,563 |
|  | PRINTS | Pfam | ProSitePatterns | ProSiteProfiles | Reactome | SFLD | SMART | SUPERFAMILY | TIGRFAM | Total |
|  | 8,518 | 37,953 | 9,970 | 22,964 | 10,299 | 91 | 19,389 | 31,616 | 1,551 | 54,176 |
|  | Blast-p/x/n hits* |  |  |  |  |  |  |  |  |  |
|  | 46,869 |  |  |  |  |  |  |  |  |  |

Additionally, the complete tables of relative gene expression obtained with *de novo* and referenced pipeline for each of the five tissues are also available in FigShare (doi: 10.6084/m9.figshare.24174285). Gene expression quantification was performed both using Fragments Per Kilobase of Transcript Per Million (FKPM) and Transcripts Per Million (TPM) metrics. The gene IDs followed the gff annotation file. The count of genes with TPM and FKPM values >1, for each tissue as identified with the two methods are displayed in Table 4. The *de novo* approach yields a significantly higher number of transcripts with TPM and FPKM values >1.

**Table 4.** Percentages of genes with Fragments Per Kilobase of transcript per Million (FKPM) and Transcripts Per Million (TPM) values >1, calculated for each tissue from the *de novo* and referenced pipeline results.

|  | Brain |  | Gills |  | Gonad |  | Liver |  | Muscle |  |
| --- | --- | --- | --- | --- | --- | --- | --- | --- | --- | --- |
|  | FKPM | TPM | FKPM | TPM | FKPM | TPM | FKPM | TPM | FKPM | TPM |
| *De novo* | 31.2 | 32.6 | 39.17 | 36.84 | 38.31 | 35.46 | 21.87 | 22.61 | 18.85 | 19.42 |
| Reference | 59.60 | 65.52 | 53.36 | 57.85 | 55.57 | 60.25 | 32.89 | 36.02 | 35.76 | 38.97 |

## Discussion

The *de novo* transcriptome assembly was obtained following a robust pipeline previously applied successfully by the team in fish as well as in other organisms such as freshwater bivalves (Machado et al. 2020; Gomes-dos-Santos et al. 2022). This is the first transcriptomic resource of *T. ligericus*, which is an important step forward considering that transcriptome assemblies are only available for a distinct *Thymallus* species, i.e., the European *T. thymallus*. Despite increasing research attention that salmonids attract, whole genome assemblies are only available for a few species (e.g. Berthelot et al. 2014; Christensen et al. 2018a,b; Varadharajan et al. 2018). This is largely due to both the large size (ca. 3 Gb) and complexity of salmonid genomes, which experienced an independent event of whole genome duplication (about 80 MY) and still show up to 25% of the genomes undergoing rediploidization (e.g. Robertson et al. 2017; Gundappa et al. 2022). These characteristics represent well-known hurdles for whole genome assembly approaches (e.g. Carruthers et al. 2018). Under this scenario, transcriptomic resources are the most suitable solution in terms of cost-effectiveness. By targeting solely transcribed regions, transcriptomes significantly reduce sequencing costs, also facilitating the computational demand and accuracy of the assembly process (e.g. Wang et al. 2019). Moreover, transcriptomes are fast, informative and valuable resources for meeting the demand for genomic tools to study threatened species (e.g. Connon et al. 2018). This is particularly important for understudied species whose conservation status is already at risk, such as many species of *Thymallus* (Weiss et al. 2021). The importance of transcriptome resources as a tool to enhance the ever-developing field of conservation physiology has been demonstrated (e.g. Connon et al. 2018) and several works have shown how transcriptome data can be used to refine species management plans (e.g. Connon et al. 2012; Narum and Campbell 2015; Wellband and Heath 2017). The present work provides a high-quality transcriptome assembly with an N50 over 1,000 bp and BUSCO scores, which are congruent with those of recently published high-quality assemblies (e.g. Lan et al. 2018; Machado et al. 2018; Moreno-Santillán et al. 2019; Li et al. 2019; Shen et al. 2020; Gomes-dos-Santos et al. 2022. BUSCO scores improved remarkably after redundancy removal with the number of duplicated genes decreasing up to 50%.

Considering the conservation status of *T. ligericus*, a vulnerable species with small and fragmented populations, the specimen capture and sacrifice must be justified. Therefore, we chose to sequence only one individual of the species. However, the robustness of the results is improved by pooling together reads belonging to five different tissues, amplifying the spectrum of represented genes. Ultimately, this approach represents a framework to develop less invasive methods for producing transcriptome resources. This approach has been previously used for other taxa such as bivalves, chondrichthyans and teleosts (Carruthers et al. 2018; Lan et al. 2018; Machado et al. 2018, 2020; Gomes-dos-Santos et al. 2022) providing extremely valuable genomic resources.

Tables of relative gene expression are presented for each of the five analyzed tissues. The analysis was carried out in parallel with two different approaches: a *de novo* pipeline and a genome-referenced pipeline yielding different values. This outcome is expected due to the differences underlying the two approaches. The referenced pipeline is based on complete genes, while the *de novo* pipeline is based on transcripts. Although the first approach is deemed as a more conservative and reliable approach, important information may be lost if the reference genome used belongs to a distinct species, as is the case herein, i.e., *T. thymallus* genome. Therefore, with the *de novo* pipeline, where we used the *T. ligericus* transcriptome here produced as a reference, we can avoid the loss of species-specific information. On the other hand, while working with transcripts through the *de novo* pipeline it is inevitable to include, for example, different unigene entries for two non-contiguous reads although they belong to the same gene. Considering the different advantages that each of the approaches brings to the analyses, both results are provided.

In conclusion, the results reported here are a step forward in the study of the vulnerable freshwater salmonid *T. ligericus*, which also represents a key resource for the genus in general. Furthermore, these resources are a valuable background for future studies inferring the vulnerability of this group to present and forthcoming threats.

## Acknowledgements

This work was supported by the Foundation for Science and Technology to GS (SFRH/BD/139069/2018 and COVID/BD/152956/2022), EF (CEECINST/00027/2021/CP2789/CT0003), LFCC (10.54499/CEECINST/00133/2018/CP1510/CT0004) and by FCT Strategic Funding UIDB/04423/2020 and UIDP/04423/2020. The authors would like to acknowledge the amateur angler Hervé Brun who caught the analyzed specimen for us in compliance with local regulations.

## Data Availability Statement

All software and respective versions applied for the present work are stated in the Methods section, together with the associated parameters. When parameters are not specified, default settings are used. Raw reads are available at NCBI Sequence Read Archive database (SRR26130661-SRR26130665; BioProject PRJNA1019285. The non-redundant transcriptome assembly is provided on NCBI databases under the accession number GKPV00000000. Redundant and non-redundant transcriptome assemblies (^*^_trinity_filtered.fasta.gz and ^*^_transcriptome.fasta.gz), transcripts (^*^_genes_final.fasta.gz), mRNA (^*^_mrna_final.fasta.gz), ORF prediction (^*^_cds_final.fasta.gz), proteins ORF prediction (^*^_proteins_final.fasta.gz), annotation files (^*^_annotation.gff3.gz) and relative gene expression results are also provided on Figshare platform (doi: 10.6084/m9.figshare.24174285).

## Authors Contributions

E. F., L. F. C. C. and S. W. conceived this work. G. S. and H. P. provided all authorizations for sampling and carried out sample collection. I. V. carried out RNA extractions. A. M. M. provided most of the scripts used for data analyses. G. S. and A. G. performed all bioinformatics analyses. G. S. and E. F. provided the first draft and all the authors read, revised, and approved the final manuscript.

## References

Andrews, S. (2010). FastQC: a quality control tool for high throughput sequence data.

Auer S, Hayes DS, Führer S, et al (2022) Effects of cold and warm thermopeaking on drift and stranding of juvenile European grayling (Thymallus thymallus). River Res Appl 401–411. 10.1002/rra.4077

Bašic T, Britton JR, Cove RJ, et al (2018) Roles of discharge and temperature in recruitment of a coldwater fish, the European grayling Thymallus thymallus, near its southern range limit. Ecol Freshw Fish 27:940–951. 10.1111/eff.12405

Bateman A, Martin MJ, O’Donovan C, et al (2017) UniProt: The universal protein knowledgebase. Nucleic Acids Res 45:D158–D169. 10.1093/nar/gkw1099

Berthelot C, Brunet F, Chalopin D, et al (2014) The rainbow trout genome provides novel insights into evolution after whole-genome duplication in vertebrates. Nat Commun 5:. 10.1038/ncomms4657

Bolger AM, Lohse M, Usadel B (2014) Trimmomatic: A flexible trimmer for Illumina sequence data. Bioinformatics 30:2114–2120. 10.1093/bioinformatics/btu170

Buchfink B, Xie C, Huson DH (2014) Fast and sensitive protein alignment using DIAMOND. Nat Methods 12:59–60. 10.1038/nmeth.3176

Camacho C, Coulouris G, Avagyan V, et al (2009) BLAST+: Architecture and applications. BMC Bioinformatics 10:1–9. 10.1186/1471-2105-10-421

Carruthers M, Yurchenko AA, Augley JJ, et al (2018) De novo transcriptome assembly, annotation and comparison of four ecological and evolutionary model salmonid fish species. BMC Genomics 19:1–17. 10.1186/s12864-017-4379-x

Christensen KA, Leong JS, Sakhrani D, et al (2018a) Chinook salmon (Oncorhynchus tshawytscha) genome and transcriptome. PLoS One 13:1–15. 10.1371/journal.pone.0195461

Christensen KA, Rondeau EB, Minkley DR, et al (2018b) The arctic charr (Salvelinus alpinus) genome and transcriptome assembly. PLoS One 13:1–30. 10.1371/journal.pone.0204076

Connon RE, D’Abronzo LS, Hostetter NJ, et al (2012) Transcription profiling in environmental diagnostics: Health assessments in Columbia River basin steelhead (Oncorhynchus mykiss). Environ Sci Technol 46:6081–6087. 10.1021/es3005128

Connon RE, Jeffries KM, Komoroske LM, et al (2018) The utility of transcriptomics in fish conservation. J Exp Biol 221:. 10.1242/jeb.148833

Costa MJ, Duarte G, Segurado P, Branco P (2021) Major threats to European freshwater fish species. Sci Total Environ 797:149105. 10.1016/j.scitotenv.2021.149105

Dainat J, Hereñú D, Pucholt P (2020) AGAT: Another Gff Analysis Toolkit to handle annotations in any GTF/GFF format. Zenodo. 10.5281/zenodo.4205393

Davidson NM, Oshlack A (2014) Corset: Enabling differential gene expression analysis for de novo assembled transcriptomes. Genome Biol 15:1–14. 10.1186/s13059-014-0410-6

Deegan LA, Golden HE, Harvey CJ, Peterson BJ (1999) Influence of Environmental Variability on the Growth of Age-0 and Adult Arctic Grayling. Trans Am Fish Soc 128:1163–1175. 10.1577/1548-8659(1999)128<1163:ioevot>2.0.co;2

Finn RD, Clements J, Eddy SR (2011) HMMER web server: Interactive sequence similarity searching. Nucleic Acids Res 39:29–37. 10.1093/nar/gkr367

Gomes-dos-Santos A, Machado AM, Castro LFC, et al (2022) The gill transcriptome of threatened European freshwater mussels. Sci Data 2022 91 9:1–10. 10.1038/s41597-022-01613-x

Grabherr MG, Haas BJ, Yassour M, et al (2011) Full-length transcriptome assembly from RNA-Seq data without a reference genome. Nat Biotechnol 29:644–652. 10.1038/nbt.1883

Gundappa MK, To TH, Grønvold L, et al (2022) Genome-Wide Reconstruction of Rediploidization Following Autopolyploidization across One Hundred Million Years of Salmonid Evolution. Mol Biol Evol 39:. 10.1093/molbev/msab310

Haas BJ, Papanicolaou A, Yassour M, et al (2013) De novo transcript sequence reconstruction from RNA-seq using the Trinity platform for reference generation and analysis. Nat Protoc 8:1494–1512. 10.1038/nprot.2013.084

Hayes DS, Lautsch E, Unfer G, et al (2021) Response of European grayling, Thymallus thymallus, to multiple stressors in hydropeaking rivers. J Environ Manage 292:. 10.1016/j.jenvman.2021.112737

Heim KC, Wipfli MS, Whitman MS, et al (2015) Seasonal cues of Arctic grayling movement in a small Arctic stream: the importance of surface water connectivity. Environ Biol Fishes 99:49–65. 10.1007/s10641-015-0453-x

Hughes NF, Reynolds JB (1994) Why do Arctic Grayling (Thymallus arcticus) get bigger as you go upstream?

Kim D, Langmead B, Salzberg SL (2015) HISAT: A fast spliced aligner with low memory requirements. Nat Methods 12:357–360. 10.1038/nmeth.3317

Labbe J, Devidal J, Albaric J (2023) Combined Impacts of Climate Change and Water Withdrawals on the Water Balance at the Watershed Scale — The Case of the Allier Alluvial Hydrosystem (France)

Lan Y, Sun J, Xu T, et al (2018) De novo transcriptome assembly and positive selection analysis of an individual deep-sea fish. BMC Genomics 19:1–9. 10.1186/s12864-018-4720-z

Langmead B, Salzberg SL (2012) Fast gapped-read alignment with Bowtie 2. Nat Methods 9:357–359. 10.1038/nmeth.1923

Li B, Dewey CN (2014) RSEM: Accurate transcript quantification from RNA-seq data with or without a reference genome. Bioinforma Impact Accurate Quantif Proteomic Genet Anal Res 41–74. 10.1201/b16589

Li B, Sun S, Zhu J, et al (2019) Transcriptome profiling and histology changes in juvenile blunt snout bream (Megalobrama amblycephala) liver tissue in response to acute thermal stress. Genomics 111:242–250. 10.1016/j.ygeno.2018.11.011

Li H, Handsaker B, Wysoker A, et al (2009) The Sequence Alignment/Map format and SAMtools. Bioinformatics 25:2078–2079. 10.1093/bioinformatics/btp352

Luecke C, MacKinnon P (2008) Landscape Effects on Growth of Age-0 Arctic Grayling in Tundra Streams. Trans Am Fish Soc 137:236–243. 10.1577/t05-039.1

Machado AM, Almeida T, Mucientes G, et al (2018) De novo assembly of the kidney and spleen transcriptomes of the cosmopolitan blue shark, Prionace glauca. Mar Genomics 37:50–53. 10.1016/j.margen.2017.11.009

Machado AM, Muñoz-Merida A, Fonseca E, et al (2020) Liver transcriptome resources of four commercially exploited teleost species. Sci Data 7:1–9. 10.1038/s41597-020-0565-9

Meyer L (2001) Spawning migration of grayling Thymallus thymallus (L., 1758) in a Northern German lowland river. Arch fur Hydrobiol 152:99–117. 10.1127/archiv-hydrobiol/152/2001/99

Moatar F, Descy J, Rodrigues S, et al (2022) The Loire River basin. 245–272

Moreno-Santillán DD, Machain-Williams C, Hernández-Montes G, Ortega J (2019) De Novo Transcriptome Assembly and Functional Annotation in Five Species of Bats. Sci Rep 9:1–12. 10.1038/s41598-019-42560-9

Mueller M, Bierschenk AM, Bierschenk BM, et al (2020) Effects of multiple stressors on the distribution of fish communities in 203 headwater streams of Rhine, Elbe and Danube. Sci Total Environ 703:134523. 10.1016/j.scitotenv.2019.134523

Narum SR, Campbell NR (2015) Transcriptomic response to heat stress among ecologically divergent populations of redband trout. BMC Genomics 16:103. 10.1186/s12864-015-1246-5

Newcombe CP, Jensen JOT (1996) Channel Suspended Sediment and Fisheries: A Synthesis for Quantitative Assessment of Risk and Impact. 693–727

Opinion AGR, De Boeck G, Rodgers EM (2020) Synergism between elevated temperature and nitrate: Impact on aerobic capacity of European grayling, Thymallus thymallus in warm, eutrophic waters. Aquat Toxicol 226:105563. 10.1016/j.aquatox.2020.105563

Pasquier J, Cabau C, Nguyen T, et al (2016) Gene evolution and gene expression after whole genome duplication in fish: The PhyloFish database. BMC Genomics 17:1–10. 10.1186/s12864-016-2709-z

Persat H, Mattersdorfer K, Charlat S, et al (2016) Genetic integrity of the European grayling (Thymallus thymallus) populations within the Vienne River drainage basin after five decades of stockings. Cybium 40:7–20

Persat H, Weiss S, Froufe E, et al (2019) A third european species of grayling (Actinopterygii, Salmonidae), endemic to the Loire river basin (France), Thymallus ligericus n. sp. Cybium 43:233–238. 10.26028/cybium/2019-433-004

Pertea M, Pertea GM, Antonescu CM, et al (2015) StringTie enables improved reconstruction of a transcriptome from RNA-seq reads. Nat Biotechnol 33:290–295. 10.1038/nbt.3122

Punta M, Coggill PC, Eberhardt RY, et al (2012) The Pfam protein families database. Nucleic Acids Res 40:290–301. 10.1093/nar/gkr1065

Quevillon E, Silventoinen V, Pillai S, et al (2005) InterProScan: Protein domains identifier. Nucleic Acids Res 33:116–120. 10.1093/nar/gki442

Richard A, Cattanéo F, Rubin JF (2015) Biotic and abiotic regulation of a low-density stream-dwelling brown trout (Salmo trutta L.) population: Effects on juvenile survival and growth. Ecol Freshw Fish 24:1–14. 10.1111/eff.12116

Robertson FM, Gundappa MK, Grammes F, et al (2017) Lineage-specific rediploidization is a mechanism to explain time-lags between genome duplication and evolutionary diversification. Genome Biol 18:1–14. 10.1186/s13059-017-1241-z

Sävilammi T, Primmer CR, Varadharajan S, et al (2019) The chromosome-level genome assembly of european grayling reveals aspects of a unique genome evolution process within salmonids. G3 Genes, Genomes, Genet 9:1283–1294. 10.1534/g3.118.200919

Shen F, Long Y, Li F, et al (2020) De novo transcriptome assembly and sex-biased gene expression in the gonads of Amur catfish (Silurus asotus). Genomics 112:2603–2614. 10.1016/j.ygeno.2020.01.026

Simão FA, Waterhouse RM, Ioannidis P, et al (2015) BUSCO: Assessing genome assembly and annotation completeness with single-copy orthologs. Bioinformatics 31:3210–3212. 10.1093/bioinformatics/btv351

Smith-Unna R, Boursnell C, Patro R, et al (2016) TransRate: Reference-free quality assessment of de novo transcriptome assemblies. Genome Res 26:1134–1144. 10.1101/gr.196469.115

Song L, Florea L (2015) Rcorrector: Efficient and accurate error correction for Illumina RNA-seq reads. Gigascience 4:1–8. 10.1186/s13742-015-0089-y

Varadharajan S, Sandve SR, Gillard GB, et al (2018) The grayling genome reveals selection on gene expression regulation after whole-genome duplication. Genome Biol Evol 10:. 10.1093/gbe/evy201

Wang B, Kumar V, Olson A, Ware D (2019) Reviving the transcriptome studies: An insight into the emergence of single-molecule transcriptome sequencing. Front Genet 10:1–11. 10.3389/fgene.2019.00384

Warren M, Dunbar MJ, Smith C (2015) River flow as a determinant of salmonid distribution and abundance: a review. Environ Biol Fishes 98:1695–1717. 10.1007/s10641-015-0376-6

Wedekind C, Evanno G, Székely T, et al (2013) Persistent Unequal Sex Ratio in a Population of Grayling (Salmonidae) and Possible Role of Temperature Increase. Conserv Biol 27:229–234. 10.1111/j.1523-1739.2012.01909.x

Weiss SJ, Gonçalves D V., Secci-Petretto G, et al (2021) Global systematic diversity, range distributions, conservation and taxonomic assessments of graylings (Teleostei: Salmonidae; Thymallus spp.). Org Divers Evol 21:25–42. 10.1007/s13127-020-00468-7

Wellband KW, Heath DD (2017) Plasticity in gene transcription explains the differential performance of two invasive fish species. Evol Appl 10:563–576. 10.1111/eva.12463

West RL, Smith MW, Barber WE, et al (1992) Autumn Migration and Overwintering of Arctic Grayling in Coastal Streams of the Arctic National Wildlife Refuge, Alaska. Trans Am Fish Soc 121:709–715. 10.1577/1548-8659(1992)121<0709:amaooa>2.3.co;2

Zuev I V., Andrushchenko PY, Chuprov SM, Zotina TA (2021) Structural Features of Scales of Baikal Grayling Thymallus baicalensis under Conditions of an Altered Hydrological Regime. Inl Water Biol 14:60–66. 10.1134/S1995082920060176

